# CD4^+^ follicular helper-like T cells are key players in anti-tumor immunity

**DOI:** 10.1101/2020.01.08.898346

**Authors:** D Singh, AP Ganesan, B Panwar, S Eschweiler, CJ Hanley, A Madrigal, C Ramírez-Suástegui, A Wang, J Clarke, O Wood, EM Garrido-Martin, SJ Chee, G Seumois, S Belanger, A Alzetani, E Woo, PS Friedmann, S Crotty, GJ Thomas, T Sanchez-Elsner, F Ay, CH Ottensmeier, P Vijayanand

## Abstract

To determine the nature of CD4^+^ T cells that provide ‘help’ for generating robust anti-tumor CD8^+^ cytotoxic T cell (CTL) responses, we profiled the transcriptomes of patient-matched CD4^+^ and CD8^+^ T cells present in the tumor micro-environment (TME) and analyzed them jointly using integrated weighted gene correlation network analysis. We found the follicular helper T cell (T_FH_) program in CD4^+^ T cells was strongly associated with proliferation and tissue-residency in CD8^+^ CTLs. Single-cell analysis demonstrated the presence of T_FH_-like cells and features linked to cytotoxic function and their provision of CD8^+^ T cell ‘help’. Tumor-infiltrating T_FH_-like cells expressed PD-1 and were enriched in tumors following checkpoint blockade, suggesting that they may respond to anti-PD-1 therapy. Adoptive transfer or induction of T_FH_ cells in mouse models resulted in augmented CD8^+^ CTL responses and impairment of tumor growth, indicating an important role of T_FH_-like CD4^+^ T cells in anti-tumor immunity.

## INTRODUCTION

CD8^+^ cytotoxic T cells (CTLs) are vital components of anti-tumor immunity (1). A distinct subset of CD8^+^ CTLs, tissue-resident memory (T_RM_) T cells, have recently emerged as critical players in mediating robust anti-tumor immune responses (2–6). Cancer immunotherapies designed to potentiate CD8^+^ CTL responses have led to remarkable clinical success, albeit in a small proportion of patients (7, 8). Therapeutic failure is due, at least in part, to an incomplete understanding of the signals and cell types in the tumor microenvironment (TME) and how they modulate effector and resident-memory CTL responses.

CD4^+^ T helper cells (T_H_) are central orchestrators of an efficient immune response. During immunization or infections, they are necessary for robust primary CD8^+^ CTL effector responses and memory transition (9–12). CD4^+^ T_H_ cells work through a multitude of mechanisms involving dendritic cell (DC) licensing and activation to initiate CD8^+^ CTL responses (13, 14) and enhance CTL proliferation (15). They mediate CD8^+^ CTL recruitment to cognate DC (16) or pathological tissue (17), and by regulation of CD8^+^ T cell TRAIL expression, they promote CTL secondary expansion on restimulation following vaccine or viral infections (18). However, within the TME, it is unknown what type of CD4^+^ T_H_ cells or CD4 helper-derived signals are essential to generate robust CD8^+^ CTL effector and T_RM_ anti-tumor immune responses.

Single-cell sequencing studies in different tumor types, have revealed tumor cell (19) or stromal cell programs (20, 21) that regulate immune response (19) or metastases (21). The analysis of tumor-infiltrating immune cells has demonstrated ‘pre-exhausted’ and exhausted CD8^+^ T cell states and multiple CD4^+^ T cell subtypes (22–24). Previous reports on CD4^+^ T cells have focused on the evaluation of specific CD4^+^ T cell subsets such as regulatory T cells (Treg), CD4^+^ T_H_1 and CD4^+^ T_H_17 cells (22, 25–27). Whilst these studies provided valuable insights into tumor-infiltrating CD8^+^ T cell or CD4^+^ T cell subsets in isolation, the molecular characterization of the cross-talk between them has not been elucidated. In order to understand the interplay between CD4^+^ T cells and CD8^+^ CTLs within the TME and the resultant impact on anti-tumor immune responses, it is critical to undertake an integrated assessment of their transcriptional programs.

We have previously reported on the transcriptomic features of tumor-infiltrating CD8^+^ CTLs in a well-characterized cohort of patients with non-small cell lung cancer (NSCLC) (3). Here, we generated the transcriptional profiles of patient-matched, purified tumor-infiltrating CD4^+^ T cells from the same cohort of patients to define the molecular interactions between CD4^+^ T cells and CD8^+^ CTLs in the TME. We performed integrated weighted gene correlation network analysis (iWGCNA) to fully characterize the molecular landscape of adaptive immune responses in the TME and the differences therein between tumors with or without robust CD8^+^ CTL effector and T_RM_ responses. We found that a follicular helper T cell (T_FH_) program in CD4^+^ T cells was strongly associated with CTL proliferation and tissue-residence in the TME. Single-cell transcriptomic analysis of tumor-infiltrating CD4^+^ T cells confirmed the presence of *CXCL13*-expressing T_FH_-like cells, which, despite expressing *PDCD1,* were enriched for transcripts linked to cell proliferation, cytotoxicity and provision of ‘help’ to CD8^+^ T cells, indicative of superior functionality. Using a murine tumor model, we showed that adoptively transferred or induced T_FH_ cells augmented CD8^+^ CTL responses and impaired tumor growth *in vivo*. Thus, based on the molecular identity and functional properties of tumor-infiltrating CD4^+^ T cells, we show that the T_FH_ subset is associated with robust anti-tumor CD8^+^ CTL and T_RM_ responses.

## RESULTS

### Follicular program in CD4^+^ T cells is associated with CD8^+^ CTL proliferation, cytotoxicity and tissue residency in the TME

To characterize the molecular interplay between cells in the TME and to define the properties in CD4^+^ T cells that are strongly associated with robust anti-tumor CD8^+^ CTL effector and T_RM_ responses across our cohort of patients with lung cancer, we developed a novel method, iWGCNA, for joint transcriptome analysis of co-expression patterns across different cell types (**Figure 1A**). Standard co-expression analysis (WGCNA (28)), is designed for studying one cell type at a time, and genes with similar expression patterns across a set of samples are grouped together into discrete clusters or modules. These modules can then be correlated with specific molecular traits to determine the group of genes whose expression is strongly associated with that trait(28). However, for a complex system, as found in the TME, where multiple cell types interact and co-regulate the trait of interest, it is important to analyze the cell transcriptomes in an integrated fashion. The goal of iWGCNA is to group transcript expression from different cell types in the TME of matched patients to form integrated gene modules that can reveal the molecular cross-talk between cell types and their relationship to specific traits.

**Figure 1.**
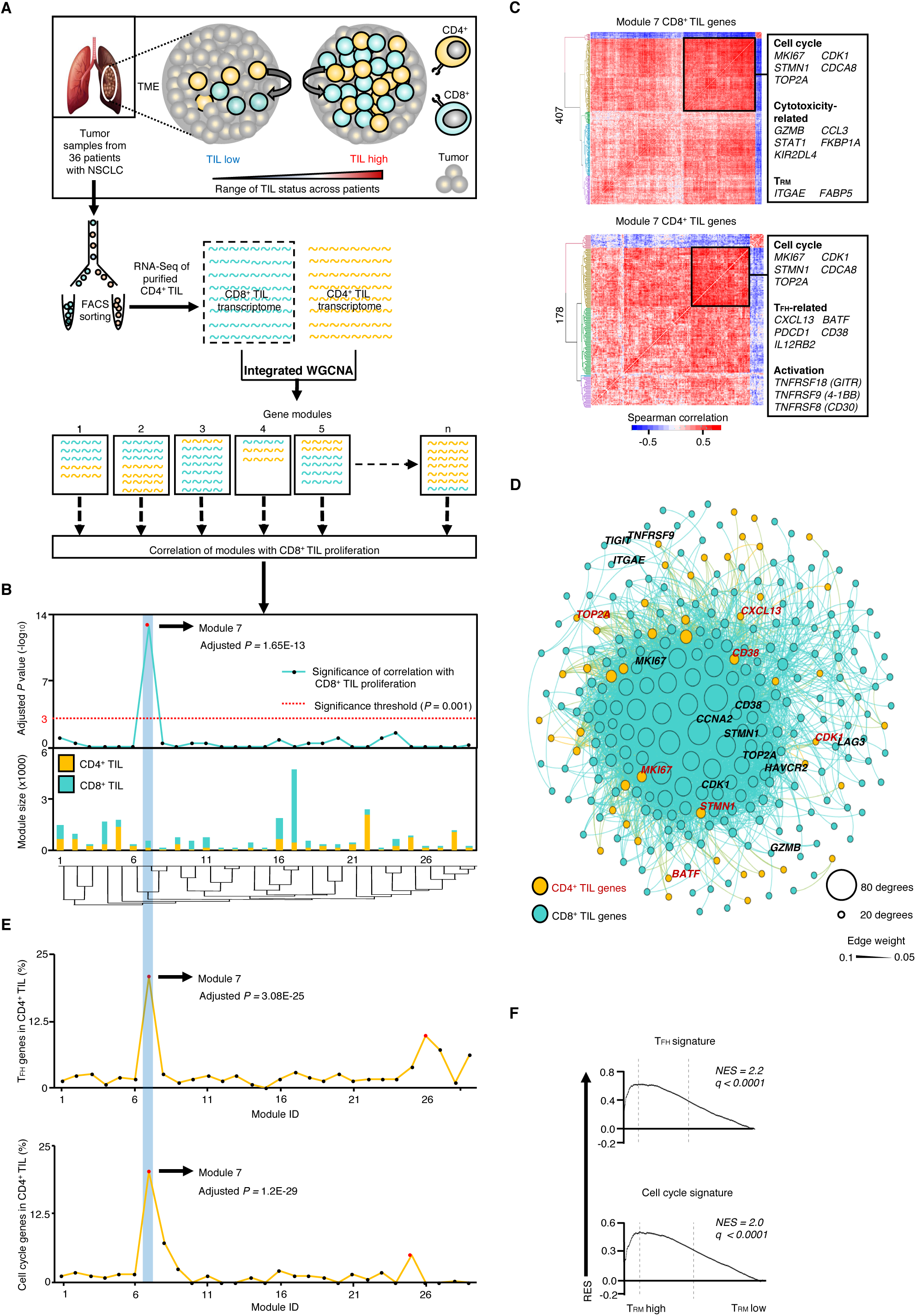
iWGCNA of transcriptomes from patient-matched CD4^+^ and CD8^+^ T cells present within lung tumors. (**A**) Schematic representation of study method, iWGCNA of the CD4^+^ and CD8^+^ TIL transcriptomes and generation of integrated modules; dotted black box indicates previously published CD8^+^ TIL transcriptomic data (3). (**B**) Barplots (bottom panel) show module size (number of genes, left margin) and composition of CD4^+^ T cell- and CD8^+^ T cell-transcripts (key above plot) for each module; number below bar represents the corresponding module ID. Significance of correlation (top panel) of proliferation-related eigengene to CD8^+^ T cell-transcripts within each module represented by symbols (Spearman correlation, left margin); red line denotes significance threshold of Bonferroni adjusted *P* value = 0.001; red symbol denotes module with significant correlation. Blue box highlights Module 7. (C) Hierarchical clustering analysis showing Spearman correlation co-expression matrix of the CD8^+^ T cell-(above) and CD4^+^-T cell transcripts (below) in Module 7 (bottom key); black frame within matrix delineates gene clusters; left margin, number of genes in module; right margin, key genes enriched in the clusters. (**D**) Module 7 genes visualized in Gephi, the nodes are colored according to cell of origin and sized according to the number of edges (connections), and the edge thickness is proportional to the strength of co-expression. (**E**) Enrichment of T_FH_ signature genes (above) or cell cycle genes (below) in CD4^+^ TILs within each module; horizontal axis represents module ID; vertical axis represents percentage of genes (symbols); red symbols denote Bonferroni adjusted *P* value < 0.001 (hypergeometric test). Blue box highlights Module 7. (**F**) GSEA of T_FH_ (top) or cell cycle signature (bottom) in the transcriptome of CD4^+^ TILs from T_RM_^high^ (*n* = 11) *versus* T_RM_^low^ (*n* = 11) tumors, presented as running enrichment score (RES) for the gene set, from most over-represented genes at left to most under-represented at right; values above the plot represent the normalized enrichment score (NES) and false discovery rate (FDR)-corrected significance value; Kolmogorov-Smirnov test.

Here, we performed iWGCNA by merging the transcriptomes from patient-matched CD4^+^ and CD8^+^ T cells present in the TME (*n* = 36) (3) and generated 29 gene network modules, each of which was composed of varying proportions of CD4^+^ T cell- and CD8^+^ T cell-transcripts (**Figure 1A** and **B**; **Supplementary Tables 1** and **2**). To determine what properties in CD4^+^ T cells were associated with robust CD8^+^ T cell responses in the TME, we correlated these gene modules with CD8^+^ T cell proliferation signature as a trait (Methods), as it represented a feature of robust anti-tumor T cell responses in the TME. Module 7 (407 CD8^+^ T cell- and 178 CD4^+^ T cell-transcripts) exhibited the highest correlation with the CD8^+^ T cell proliferation signature (*r* = 0.915, adjusted *P* = 1.65E-13) (**Figure 1B**) and, as expected, nearly 25% of the CD8^+^ T cell-transcripts in Module 7 were cell cycle-related genes. Clustering analysis of the 407 CD8^+^ T cell-transcripts present in Module 7 identified a tightly correlated and co-expressed subset of transcripts (*n* = 171), which, besides cell cycle genes, included several genes encoding products linked to effector and cytotoxic functions such as *GZMB, CCL3, STAT1, FKBP1A, KIR2DL4* (**Figure 1C** and **1D**; **Supplementary Table 2**). This highly co-expressed cluster of CD8^+^ T cell-transcripts also contained *ITGAE* (which encodes for the α−subunit of the integrin molecule α_E_β_7_), a well-established marker of lung CD8^+^ T_RM_ cells (29, 30), which we have recently shown to be a critical determinant of survival outcomes in lung cancer (**Figure 1C** and **1D**; **Supplementary Table 2**) (3). Together, these findings indicated that cell proliferation, effector functions and T_RM_ features are highly correlated and interconnected processes in CD8^+^ CTLs present in the TME.

To assess the properties in CD4^+^ T cells that are associated with these functional features of CD8^+^ CTLs in the TME, we next analyzed the CD4^+^ T cell-transcripts (*n* = 178) present in Module 7. We observed a tightly correlated and co-expressed cluster of transcripts (*n* = 61), which included several genes linked to follicular helper CD4^+^ T cells (T_FH_), such as *CXCL13, BATF, CD38*, *PDCD1* (31) (**Figure 1C** and **1D**; **Supplementary Table 2**). T_FH_ cells are the principal CD4^+^ T cell subpopulation that provides essential ‘help’ to B cells, promoting maturation of antibody affinity in germinal centers (GC) (32). Among the T_FH_-related transcripts in this cluster, *BATF* encodes for a transcription factor involved in T_FH_ differentiation (33). CXCL13 is a chemokine produced by human T_FH_ cells, but not by their murine counterparts; CXCL13 has been shown to play an important role in the homing of B cells to follicles (34–36). The Module 7 cluster was also composed of CD4^+^ T cell-transcripts linked to cell proliferation (e.g., *KI67, TOP2A, STMN1, CDK1)* and T cell activation, such as *CD38* (37–39), *TNFRSF9* (which encodes for 4-1BB (40, 41)), *TNFRSF18* (which encodes for GITR (42)), and *TNFRSF8* (which encodes for CD30 (43)) (**Figure 1C**; **Supplementary Table 2**). In an unbiased overlap analysis using hypergeometric test, we confirmed that among the 29 gene modules generated from the iWGCNA, module 7 exhibited the highest enrichment for both the T_FH_ and cell cycle signature genes (**Figure 1E**). Consistent with these findings, GSEA also showed significant enrichment of proliferation and T_FH_ gene signatures in tumor-infiltrating CD4^+^ T cells from tumors with high T_RM_ density relative to those with low T_RM_ density (**Figure 1F**). These results suggest that the T_FH_ program in CD4^+^ T cells is tightly coupled with features of cell proliferation and activation in the TME. Taken together with results from the CD8^+^ T cell-transcripts in Module 7 (**Figure 1C**), our iWGCNA analysis indicates that a T_FH_-like transcriptional program in CD4^+^ T cells is strongly associated with CD8^+^ CTL proliferation, effector function and T_RM_ features in the TME, all features of a robust anti-tumor immune response.

### Major transcriptional changes characterize tumor-infiltrating T_FH_-like cells

T_FH_ cells have been reported within human cancers, where they were shown to support B cell responses, tertiary lymphoid structures and were associated with improved survival outcomes (44, 45). Recent studies in murine breast cancer models showed the importance of IL21, presumed to be T_FH_-derived, in mediating response to checkpoint blockade by impacting B cell infiltration and antibody production (46). Given that we found association of tumor-infiltrating T_FH_-like cells with CD8^+^ CTL responses, we next sought to characterize these T_FH_ cells in order to underpin their mechanistic role in anti-tumor immunity.

Conventional GC T_FH_ cells are characterized by the expression of CXCR5, however in the context of tumor and inflammation, T_FH_ cells lacking expression of CXCR5 have also been described (47, 48). Therefore, to capture the entire spectrum of CD4^+^ T cells with a follicular program, we performed single-cell RNA-seq of both CXCR5^+^ and CXCR5^−^ CD4^+^ T cells in the TME (**Figure 2A**; **Supplementary Table 1**). Unbiased single-cell transcriptomic analysis of 5317 tumor-infiltrating CD4^+^ T cells revealed 9 distinct clusters (**Figure 2B**). Given that GC T_FH_ are the major producers of the B cell chemoattractant CXCL13, we utilized *CXCL13* expression to mark CD4^+^ T cells with a follicular program. Cells expressing *CXCL13* transcripts were highly enriched in cluster 3 (∼70% of cells expressed *CXCL13*), which suggests that *CXCL13*-expressing cells likely represent a distinct CD4^+^ T cell subset (**Figure 2B**; **Supplementary Figure 1A** and **1B**; **Supplementary Table 3**). Single-cell analysis of the 2783 lung-infiltrating CD4^+^ T cells (N-TIL) showed very little *CXCL13* expression (**Supplementary Figure 1C**). These findings were confirmed at the protein level by flow cytometric analysis of CXCL13 expression in tumor and lung tissue (**Supplementary Figure 1D**).

**Figure 2.**
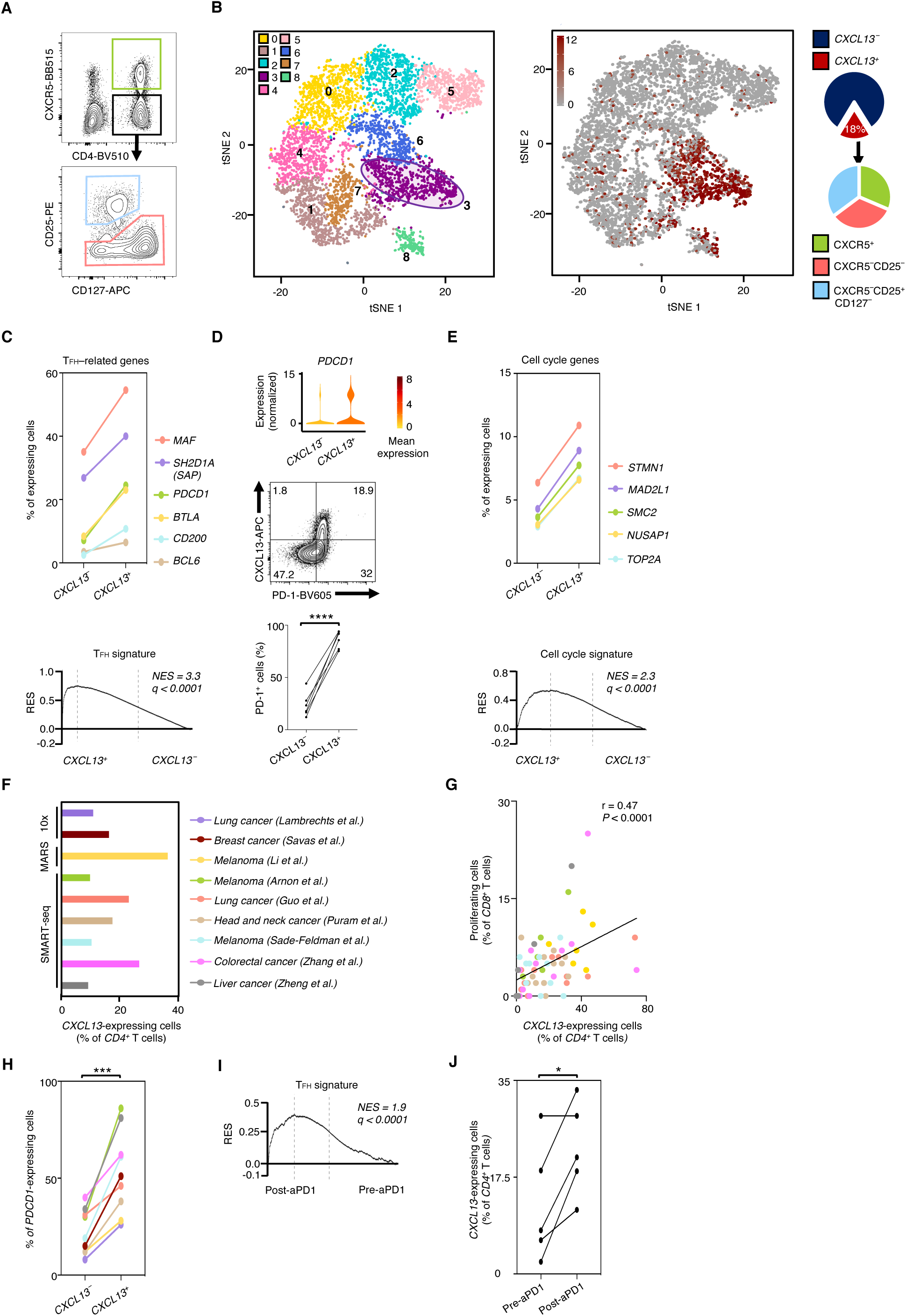
Single-cell transcriptomics reveal molecular profile of *CXCL13*-expressing tumor-infiltrating CD4^+^ T cells. (**A**) Sorting strategy for single-cell RNA-seq assays. Live, singlet gated, CD14^−^ CD19^−^CD20^−^CD8^−^CD56^−^CD45^+^CD3^+^CD4^+^ lymphocytes from 6 lung tumors were sorted as CXCR5^+^ (green), CXCR5^−^CD25^+^CD127^−^ (blue) and CXCR5^−^CD25^−^ (coral) subsets. (**B**) Seurat clustering of 5317 single cell TIL transcriptomes identifying 9 clusters (left); each symbol represents a cell; circle delineates *CXCL13* cluster. tSNE visualization of cells in **B** (right); each symbol represents a cell; brown color indicates *CXCL13* expression in counts per million (CPM). Pie chart represents the percentage of *CXCL13*-expressing cells (*n* = 955) among all TILs (far right, above) and relative proportions of each of the sorted subsets that express *CXCL13* (far right, below). (**C**) Percentage (left margin) of *CXCL13*-non-expressing or *CXCL13*-expressing cells that express the indicated T_FH_-related genes. Below, GSEA of T_FH_ signature in the transcriptome of *CXCL13*-expressing *versus* CXCL13-non-expressing cells, presented as in Figure 1F. (**D**) Violin plots (above) of expression of *PDCD1* in *CXCL13*-non-expressing or *CXCL13*-expressing cells; shape represents the distribution of expression among cells and color represents expression (log_2_(CPM+1)). Flow-cytometric analysis (middle) shows the expression of PD-1 and CXCL13 in live, singlet-gated, CD45^+^CD3^+^CD4^+^ TILs (*n* = 6) from patients with NSCLC. Plot (below) shows percentage of PD-1^+^ cells in CD4^+^CXCL13^-^ or CD4^+^CXCL13^+^ TILs; **** *P <* 0.0001 (two-tailed paired Student’s *t*-test). (**E**) Percentage (left margin) of *CXCL13*-non-expressing or *CXCL13*-expressing cells that express the indicated cell cycles genes. Below, GSEA of cell cycle signature in the transcriptome of *CXCL13*-expressing *versus* CXCL13-non-expressing cells, presented as in Figure 1F. (**F**) Percentage of *CXCL13*-expressing cells (horizontal axis) in tumor-infiltrating *CD4^+^* T cells derived from integrated analysis of 9 single-cell RNA-seq datasets (right, key). (**G**) Correlation of frequencies of patient-matched *CXCL13*-expressing *CD4^+^* TILs with cell cycle genes-expressing *CD8^+^* TILs in the assessed datasets (*n* = 63) (key as in **F**); each dot represents a donor; *r* value indicates the Spearman correlation coefficient. (**H**) Percentage (left margin) of *CXCL13*-non-expressing or *CXCL13*-expressing *CD4^+^* TILs that express *PDCD1* transcripts in the assessed datasets (key as in **F**); *** *P <* 0.0005 (two-tailed paired Student’s *t*-test). (**i**) GSEA of T_FH_ signature in the transcriptome of *CD4^+^* TILs from pre-*versus* post-therapy tumor samples obtained from patients treated with aPD-1 mAB, presented as in Figure 1F. (**J**) Percentage (left margin) of *CXCL13*-expressing cells in *CD4^+^* TILs from matched pre- or post-therapy tumor samples obtained from patients (*n* = 5) treated with aPD-1 mAB; * *P <* 0.05 (two-tailed paired Student’s *t*-test).

Single-cell differential gene expression analysis of the *CXCL13*-expressing *versus CXCL13*-non-expressing CD4^+^ tumor-infiltrating lymphocytes (TILs) (Methods) revealed over 1000 differentially expressed transcripts (**Supplementary Table 4**). We found both higher expression and higher percentage of cells expressing T_FH_-related genes (*MAF, SH2D1A, PDCD1, BTLA, CD200, BCL6*(*31*)) in *CXCL13*-expressing cells than in *CXCL13*-non-expressing cells (**Figure 2C**; **Supplementary Figure 1E**). *PDCD1* encodes for the inhibitory checkpoint molecule PD-1, which is known to be constitutively expressed in GC T_FH_ cells and regulates GC localization and helper functions (49). Similarly, the majority of CXCL13^+^CD4^+^ TILs expressed PD-1 when compared to CD4^+^ TILs not expressing CXCL13 (**Figure 2D**). GSEA confirmed significant enrichment of T_FH_ signature genes in the *CXCL13*-expressing cells (**Figure 2C**). Together, these findings clearly established that the expression of *CXCL13* delineated a T_FH_ program, *i.e., CXCL13*-expressing cells represent T_FH_-like cells in TME. We found that *CXCL13*-expressing cells were present in equal proportions in CXCR5^+^ and the two CXCR5^−^ subsets (CD25^+^CD127^−^ and CD25^−^) (**Figure 2B**; **Supplementary Figure 1B**), which indicated that using CXCR5 as a surface marker for T_FH_ cells would have missed the majority of T_FH_ cells. Consistent with iWGCNA results (**Figure 1C-E**), higher proportions of *CXCL13*-expressing cells expressed cell cycle-related transcripts, and GSEA also demonstrated enrichment of cell cycle genes in the *CXCL13*-expressing cells (**Figure 2E**). Together these results indicate that despite expressing high levels of *PDCD1* (**Figure 2C** and **D**), T_FH_-like cells actively proliferate in the tumor microenvironment presumably in response to tumor-associated antigens and may be important cellular targets of anti-PD-1 therapies.

In agreement with our results, analysis of nine published single-cell studies of CD4^+^ TIL transcriptomes (*n* = 25,149) from several cancer types showed that *CXCL13*-expressing T_FH_-like cells represented 9-36% of the CD4^+^ T cells in the TME (**Figure 2F**; **Supplementary Table 5**). Importantly, we found a positive correlation between the proportion of *CXCL13*-expressing T_FH_-like CD4^+^ T cells and proliferating CD8^+^ T cells in the TME (*n* = 63, **Figure 2G**). Across all cancer types, a greater proportion of *CXCL13*-expressing T_FH_-like cells expressed *PDCD1* transcripts relative to their *CXCL13*-non-expressing counterparts (**Figure 2H**). Hence, we evaluated whether anti-PD-1 therapies targeted T_FH_-like CD4^+^ T cells, which, if so, would lead to their enrichment in tumor samples post-treatment with anti-PD-1 agents. As expected, we found a strong enrichment of T_FH_ signature genes in CD4^+^ TIL transcriptomes and an increase in the proportion of *CXCL13*-expressing T_FH_-like cells from post-treatment samples compared to pre-treatment samples (**Figure 2I** and **J**). Overall, these results indicate that T_FH_-like cells are an important CD4^+^ T cell subset in the TME that is linked to robust CD8^+^ CTL responses, and they are likely to be responsive to anti-PD-1 therapy.

### Tumor-infiltrating T_FH_-like CD4^+^ T cells possess superior functional properties

To further probe the functional properties of these T_FH_-like cells, we performed pathway analysis of the transcripts differentially expressed in *CXCL13*-expressing cells relative to *CXCL13*-non-expressing cells in our single-cell CD4^+^ TIL transcriptomes. *CXCL13*-expressing T_FH_-like cells showed significant enrichment of pathways linked to co-stimulation (CD28 and ICOS-ICOSL signaling), which is important for T_FH_ activation (31) (**Figure 3A**; **Supplementary Table 6**). As expected of activated and co-stimulated T cells in the tumor, these T_FH_-like cells showed increased expression of transcripts encoding for cytokines (IFN-γ and IL21) and co-stimulation molecules (GITR and OX-40), which are known to play an important role in CD4^+^ T cell-mediated ‘help’ to CD8^+^ CTLs (50–56) (**Figure 3B**). Notably, the cytokine IL21 has been shown to support CD8^+^ CTL survival and function (50–53). Another important finding from our analysis was that Tregs, were seen mainly within CXCL13-non-expressing CD4^+^ T cells, which also showed differential expression of *FOXP3* transcripts (**Supplementary Figure 2A**).

**Figure 3.**
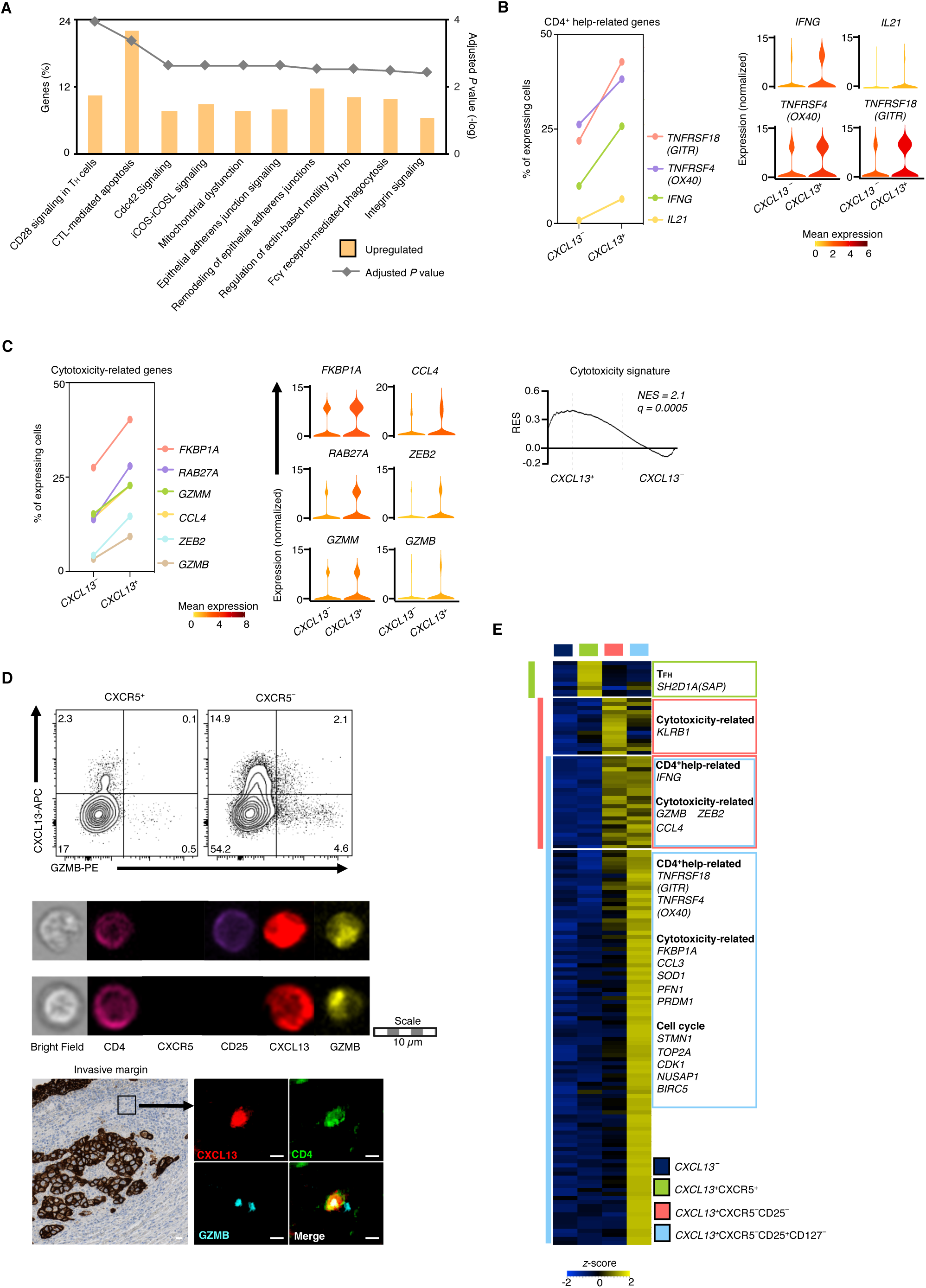
Highly functional T_FH_-like CD4^+^ T cells were CXCR5 negative. (**A**) Canonical pathways (horizontal axis; bars in plot) for which *CXCL13*-expressing TILs show enrichment, presented as the frequency of differentially expressed genes encoding components of each pathway that are upregulated (key) in *CXCL13*-expressing TILs relative to their expression in *CXCL13*-non-expressing TILs (left vertical axis), and adjusted *P* values (right vertical axis; grey squares; Benjamini-Hochberg test); *P* < 0.05. (**B**) Percentage (left margin) of *CXCL13*-non-expressing or *CXCL13*-expressing cells that express the indicated CD4^+^ help-related genes (left plot) and violin plots of expression of the same genes in *CXCL13*-non-expressing or *CXCL13*-expressing cells (right), presented as in Figure 2D. (**C**) Percentage (left margin) of *CXCL13*-non-expressing or *CXCL13*-expressing cells that express the indicated cytotoxicity-related genes (left plot) and violin plots (middle) of expression of the same genes in *CXCL13*-non-expressing or *CXCL13*-expressing cells, presented as in Figure 2D. GSEA (right) of cytotoxicity signature in the transcriptome of *CXCL13*-expressing *versus* CXCL13-non-expressing cells, presented as in Figure 1F. (**D**) Flow-cytometric analysis (top panel) shows expression of CXCL13 and granzyme B in CXCR5^+^ and CXCR5^−^ subsets in live, singlet-gated, CD45^+^CD3^+^CD4^+^ TILs (*n* = 9) from patients with NSCLC; numbers in quadrants indicate percentage of CD4^+^ TILs in each. ImageStream analysis (middle panel) shows expression of CXCL13 and granzyme B in live, singlet-gated, CD3^+^CD4^+^CXCR5^−^ TILs (*n* = 3). Pan-CytoKeratin(CK) (DAB) IHC staining (bottom panel, left) shows invasive margin (black frame) of tumors (*n* = 41); scale bar represent 200µm. PseudoIF image of MxIHC staining (bottom panel, right) shows CXCL13 (red), CD4 (green), granzyme B (cyan) in region indicated by arrow; scale bars represent 10µm. (**E**) Expression of transcripts differentially expressed in *CXCL13*-expressing *versus CXCL13*-non-expressing TILs, in various sorted subsets (above heatmap, right key). Each column represents the average expression (CPM) in a particular subset. Left margin, vertical colored lines indicate subset in which the genes are differentially expressed. Right margin, examples of key transcripts expressed uniquely or shared by corresponding subsets.

A surprising finding was enrichment of the cytotoxicity pathway in *CXCL13*-expressing cells (**Figure 3A** and **3C**; **Supplementary Table 6**). We found higher expression and a higher percentage of cells expressing cytotoxicity-related transcripts (57) such as *GZMB, GZMM, FKBP1A, RAB27A, CCL4* and *ZEB2* in *CXCL13*-expressing than in *CXCL13*-non-expressing cells (**Figure 3C**). This finding was confirmed by GSEA, which showed significant enrichment of cytotoxicity signature genes in the *CXCL13*-expressing cells (**Figure 3C**; **Supplementary Figure 2B**). We verified the expression of granzyme B (a canonical cytotoxicity marker) in tumor-infiltrating CD4^+^ T_FH_-like cells (CXCL13-expressing CD4^+^ T cells) using three independent approaches: a) intracellular staining by flow cytometry, b) ImageStream imaging cytometry, and c) immunohistochemical (IHC) analyses of human lung tumor samples (**Figure 3D**).

Because anti-tumor functions were not previously ascribed to conventional T_FH_ cells, we sought to investigate the precise nature of the cells that harbored these functions using our single-cell RNA-seq data where cells are annotated based on CXCR5 protein expression (**Figure 2A**). Since the *CXCL13*-expressing T_FH_-like cells were present in equal proportions in CXCR5^+^ and the two CXCR5^−^ subsets (CD25^+^CD127^−^ and CD25^−^), we first asked whether the superior functional properties were attributes of all or were unique to one subset. Pairwise comparisons for differential expression analysis between the three subsets showed that T_FH_ signature genes were significantly enriched in all three T_FH_ subsets (**Supplementary Figure 2C**), which suggested that all subsets had switched on a T_FH_ molecular program. However, transcripts linked to superior anti-tumor properties, such as cytotoxicity (*KLRB1*, *GZMB, CCL3, CCL4, FKBP1A, SOD1, ZEB2* (*57*)) and provision of CD8^+^ T cell ‘help’ (*IFNG*, *GITR*, *OX40*) were mainly enriched in both CXCR5^−^ T_FH_ subsets (**Figure 3E**; **Supplementary Figure 2C**; **Supplementary Table 7**). Notably, cell cycle-related transcripts were also expressed predominantly in this CXCR5^−^ T_FH_ subset, which together suggest that the functionally important T_FH_ cells that proliferate in the TME are contained within this subset (**Figure 3E**; **Supplementary Figure 2C**; **Supplementary Table 7**).

### T_FH_-like CD4^+^ T cells colocalize with CD8^+^ T_RM_ cells in the TME

We undertook multi-parametric immunohistochemistry to gain insights into the organization of CD4^+^ T_FH_-like cells within the TME, and importantly, determine the spatial relationship between tumor cells, tertiary lymphoid structures (TLS) and CD8^+^ T_RM_ cells, the density of which has been linked to good survival outcomes (2–6). As expected, CXCL13-expressing T_FH_-like CD4^+^ T cells were localized in TLS (48), but they were also present in the tumor core and its invasive margins (**Figure 4A**). T_RM_ cells (CD8^+^CD103^+^ cells) in the tumor core and invasive margins were seen in close proximity to CXCL13-expressing CD4^+^ T cells, which suggested potential for crosstalk and ‘help’ (**Figure 4B**). Therefore, we asked whether the density of T_FH_-like CD4^+^ T cells in the tumor core and invasive margins was positively associated with density of T_RM_ cells in the tumor. We found a significant positive correlation (*r* = 0.72, *P* < 0.0001) between the density of T_RM_ cells (CD8^+^CD103^+^ cells) and T_FH_ cells (CD4^+^CXCL13^+^ cells) in both tumor core and invasive margins (**Figure 4C**). Importantly, the proportion of CD4^+^ T cells that were CXCL13^+^ also positively correlated (*r* = 0.58, *P* < 0.0001) with the density of T_RM_ cells (CD8^+^CD103^+^ cells) (**Figure 4C**), a finding that further supports the association between a follicular program in CD4^+^ T cells and robust CD8^+^ T cell responses in tumors.

**Figure 4.**
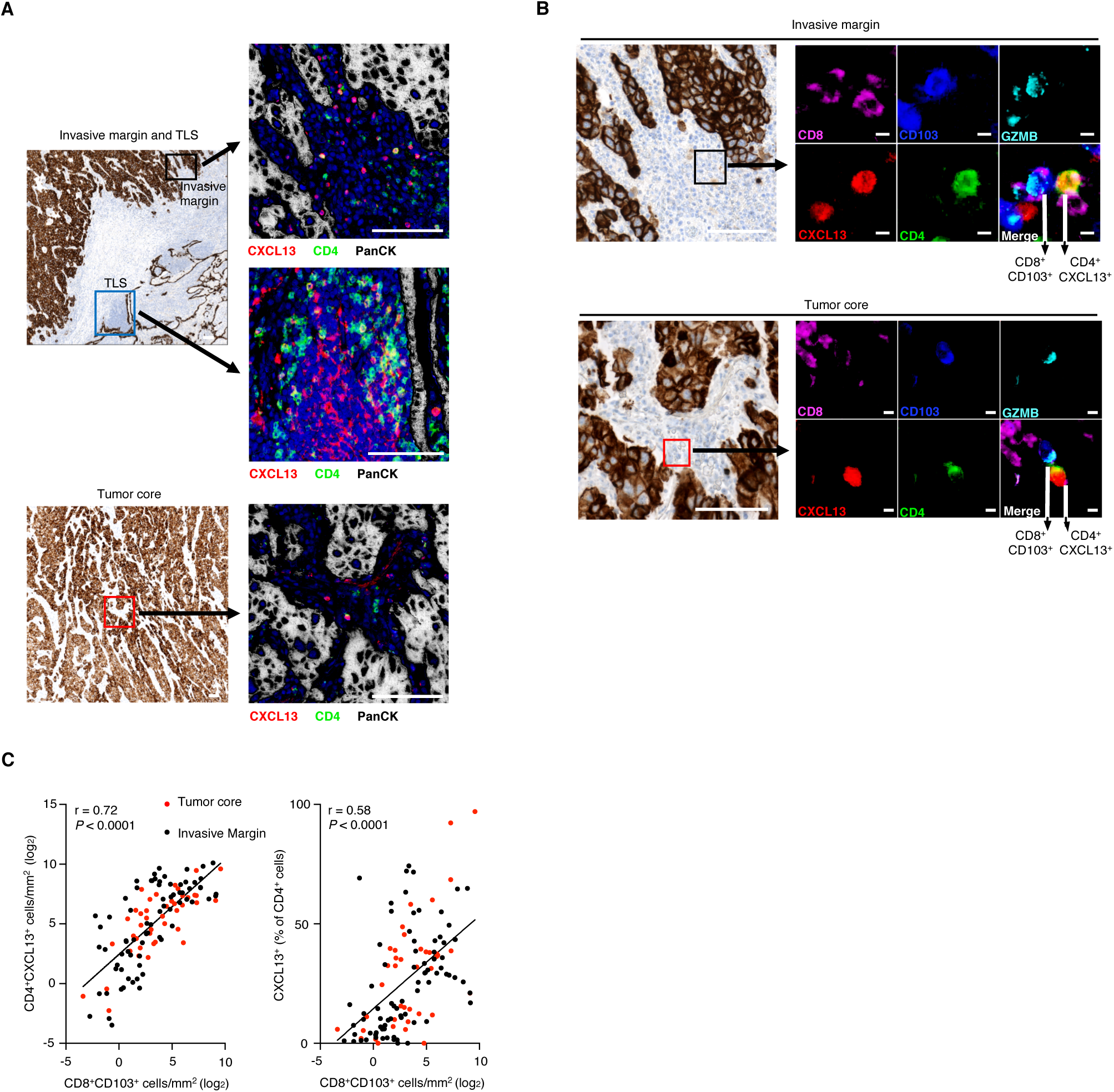
T_FH_-like cells infiltrate tumor and associate with CD8^+^ T_RM_ cells. (**A**) Pan-CK (DAB) IHC staining shows TLS (blue frame), invasive margin (black frame)(above, left image) and tumor core (red frame)(below, left image). PseudoIF image of MxIHC staining shows pan-CK (white), CD4 (green), CXCL13 (red), nuclei (blue) in regions indicated by arrows; scale bars represent 200µm. (**B**) Pan-CK (DAB) IHC staining shows invasive margin (black frame)(above, left image) and tumor core (red frame)(below, left image); scale bars represent 200µm. PseudoIF image of MxIHC staining shows CD8 (magenta), CD103 (blue), granzyme B (cyan), CXCL13 (red), CD4 (green) in regions indicated by arrows; scale bars represent 10µm. (**C**) Correlation of the number of CD4^+^CXCL13^+^ cells and CD8^+^CD103^+^ cells in lung tumors (quantified by IHC)(left). Correlation of the percentage of CXCL13^+^ cells in CD4^+^ cells and the number of CD8^+^CD103^+^ cells in lung tumors (quantified by IHC)(right). Each symbol (key) represents an image; 3 images analyzed per patient (*n* = 41); *r* values indicate the Spearman correlation coefficient; *P* values, Spearman correlation.

### T_FH_ cells augment CD8^+^ CTL responses and impair tumor growth

In order to evaluate the functional role of antigen-specific T_FH_ cells in anti-tumor immunity *in vivo*, we utilized the B16F10-OVA murine syngeneic tumor model in which the aggressive growth of melanoma tumors is unhindered by most interventions (58). OT-II TCR transgenic (specific for OVA 323-339) mice were first immunized with OVA in alum to generate T_FH_ cells *in vivo*. We then adoptively transferred OT-II CD4^+^ T_FH_ cells or OT-II CD4^+^ T_EFF_ cells to tumor-bearing mice at 11 days post-tumor inoculation and assessed the fold increase in tumor volume over the next 48 hours (**Figure 5A**; **Supplementary Figure 3**). Only the transfer of OT-II CD4^+^ T_FH_ cells resulted in significant reduction in tumor growth when compared to mice that received no adoptive transfer (**Figure 5A**). As an alternative strategy, we adoptively transferred naïve OT-II CD4^+^ cells and immunized tumor-bearing mice to induce antigen-specific T_FH_ responses *in vivo* and assessed effects on tumor growth. Immunized mice demonstrated significant reduction in tumor volume relative to unimmunized controls (**Figure 5B**). The tumors in immunized mice had a higher proportion of T_FH_ cells (**Figure 5C**; **Supplementary Fig. 4**), and notably, the tumor-infiltrating T_FH_ cells were CXCR5^neg^ similar to the T_FH_-like cells observed in human tumors. Enhanced T_FH_ cell infiltration was also accompanied by an increased frequency of CD8^+^ T cells, higher proportions of which also expressed granzyme B and Ki-67, implying greater cytotoxic potential and cell proliferation, respectively (**Figure 5C**). Taken together, these results reveal that induction of T_FH_ response functionally bolsters anti-tumor CD8^+^ CTL response and improves tumor control.

**Figure 5.**
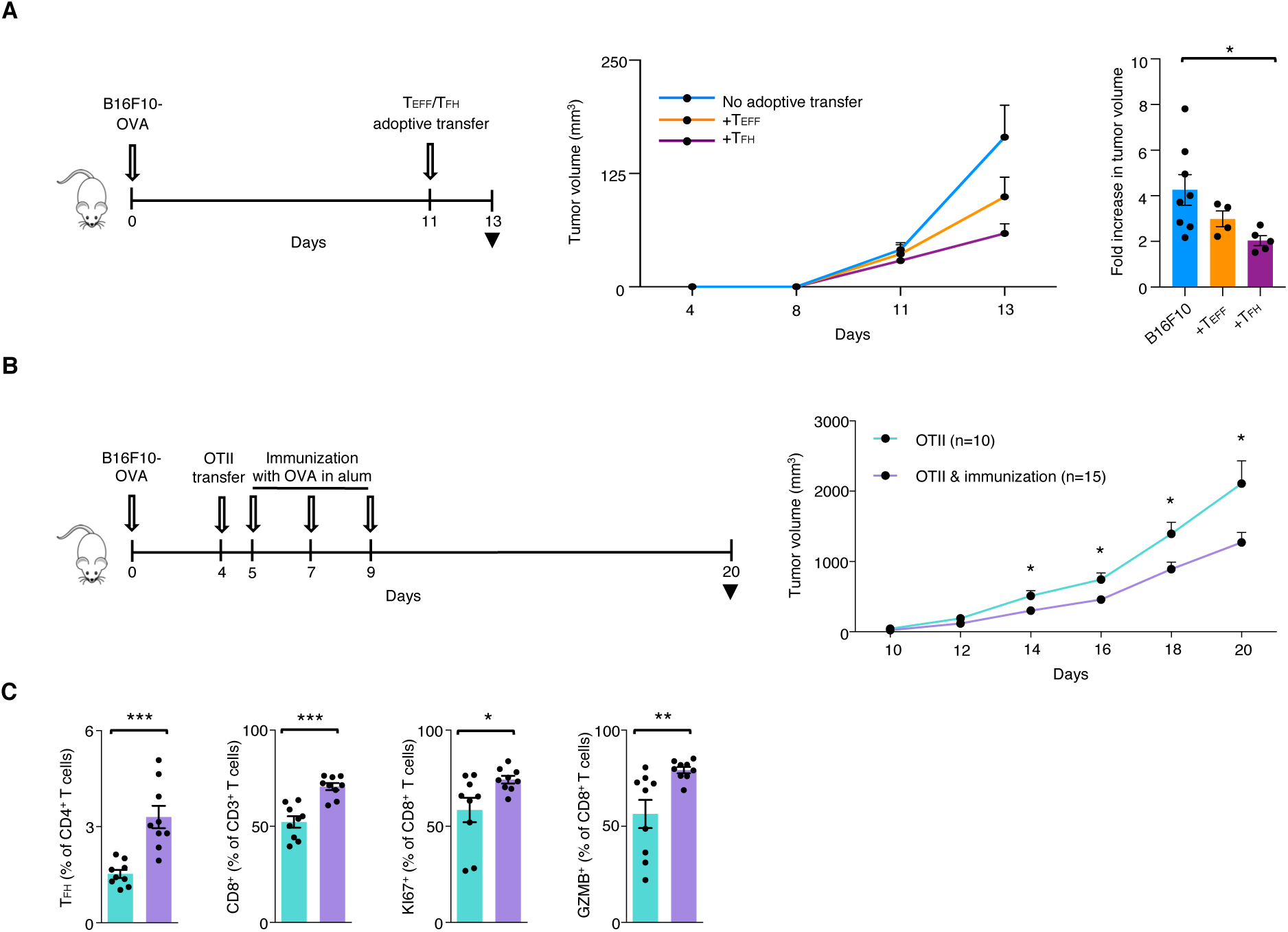
Induction of T_FH_ cells in tumor-bearing mice promotes anti-tumor immunity. (A) Mice were *s.c.* inoculated with B16F10-OVA and received adoptive transfer of OT-II CD4^+^ T_EFF_ cells or OT-II CD4^+^ T_FH_ cells at the indicated time point. Tumor volume (left) and fold increase in tumor volume (right) from day 11 to day 13 in mice (n = 4-8/group) treated as indicated (error bars are mean ± SEM); * *P <* 0.05 (Mann-Whitney test); data representative of two independent experiments. (**B,C**) Mice were *s.c.* inoculated with B16F10-OVA and received adoptive transfer of naïve OT-II CD4^+^ cells followed by immunization with OVA in alum at the indicated time points. Tumor volume (**B**) and tumor-infiltrating cell frequencies assessed by flow cytometric analysis (**C**) in mice (n = 10-15/group) treated as indicated (error bars are mean ± SEM); * *P <* 0.05, ** *P <* 0.005, *** *P <* 0.0005 (Mann-Whitney test); data representative of two independent experiments.

## DISCUSSION

Published transcriptional studies of tumor-infiltrating CD4^+^ T cells from patients with cancer have largely focused on the analysis of specific CD4^+^ T cell subtypes or CD4^+^ TILs in isolation, without integration with CD8^+^ CTL responses (25, 27, 59). We interrogated the transcriptomes of patient-matched CD4^+^ and CD8^+^ TILs using a novel iWGCNA-based approach to identify molecular features of CD4^+^ TILs that were linked to robust CD8^+^ anti-tumor immune responses and thus better patient outcomes. This combined TIL transcriptomic dataset, generated from patient-matched samples, facilitated the capture of *in vivo* TIL interactions within the TME and functional mapping of CD4^+^ TIL responses with those of CD8^+^ TILs. Such cell-specific, context-dependent, cross-talk would have otherwise been challenging to decipher in patients.

Using our cohort of patients with a range of CD8^+^ and T_RM_ TIL densities, we discovered a link between a T_FH_ program in CD4^+^ TILs and features of robust CD8^+^ T cell anti-tumor immune responses such as proliferation, cytotoxicity and tissue residency. Furthermore, we found that T_FH_-like CD4^+^ TILs possessed superior functional properties including proliferation, cytotoxic potential and provision of ‘help’ to CD8^+^ CTLs. Conventional GC T_FH_ are known to provide B cell ‘help’ during viral infections by promoting GC development, B cell affinity maturation and class switch recombination (31). However, the association of tumor-infiltrating T_FH_ cells with CD8^+^ T cell ‘help’ and robust CD8^+^ CTL responses within tumors has not been described before. We uncovered increased expression of transcripts encoding molecules that mediate CD8^+^ T cell ‘help’ (*TNFRSF18, TNFRSF4, IFNG, IL21*) in T_FH_-like CD4^+^ TILs. CD4^+^ helper-derived IL21 has a prominent role in CD8^+^ T cell ‘help’ by inducing the BATF-IRF4 axis to sustain CD8^+^ T cell maintenance and effector response (50, 51, 60). OX40 and GITR signaling on CD4^+^ T cells critically impacts CD8^+^ T cell priming, accumulation and expansion (55, 56). The role of interferon-γ produced by CD4^+^ T cells in helping CD8^+^ CTLs and CD8^+^ T_RM_ cells has been well established (54, 61). We further showed the co-localization of CD8^+^ T_RM_ cells with CD4^+^CXCL13^+^ T_FH_ cells in tumor invasive margins and tumor core, which lends support to the notion that T_FH_ cells may mediate CD8^+^ T cell ‘help’.

Our single-cell RNA-seq data unraveled another novel finding, the expression of granzyme B by the T_FH_-like CD4^+^ TILs. The existence of MHC class II-restricted CD4^+^ CTLs has been demonstrated in viral infections where they may play a particularly important role in viral clearance in the face of virus strategies to escape CD8^+^ CTL responses (62). Recently, GC T_FH_ cells with cytotoxic potential have been reported in recurrent tonsillitis (63). Hence, it is plausible to hypothesize that a CTL program may also be induced in tumor-infiltrating T_FH_ cells within tumors that have downregulated their MHC I expression. Interestingly, TGF-β signaling has been shown to promote differentiation of both CD4^+^ CTLs (64–66) and CD8^+^ T_RM_ cells (30, 67), whilst IL2 depletion may induce CXCL13 production and T_FH_ development (68). Strikingly, both these signaling cues are abundant within tumors that harbor Tregs, suggesting that such CD4^+^ T cell plasticity may have evolved as a mechanism to provide these TILs with survival fitness in such immunosuppressive IL2-deprived environments. We utilized a number of complementary methods and provided a spatially resolved analysis to confirm the presence and location of these GZMB- and CXCL13-expressing CD4^+^ TILs.

In further dissecting the molecular profile of the T_FH_-like CD4^+^ TILs at single-cell resolution, we revealed T_FH_ features in both the CXCR5^+^ and CXCR5^−^ CD4^+^ T cell subsets. Our results are consistent with recent studies both in breast cancer and rheumatoid arthritis, which demonstrated the presence of CXCL13-producing T_FH_ cells that lacked CXCR5 expression (47, 48). An additional finding from our studies was that the superior functional properties such as cytotoxicity, provision of CD8^+^ ‘help’ and proliferation observed in T_FH_-like CD4^+^ TILs, specifically resided in the CXCR5^−^ subset.

Previous studies have reported the presence of T_FH_ cells in cancers, where they were shown to impact B cell activation and antibody production (44–46). Our study presents novel insights into the functional role of T_FH_ cells in augmenting CD8^+^ CTL responses within tumors. In addition to our human data, functional evidence is provided by our *in vivo* murine studies, in which induction of T_FH_ response was associated with increased CD8^+^ CTL infiltration, proliferation and granzyme B expression within tumors, culminating in tumor control. A further important implication for T_FH_ cells derives from their high PD-1 expression, rendering them targets of anti-PD-1 therapy. CD8^+^ CTL subsets are considered to be the primary cellular responders to anti-PD-1 therapy, however, our re-analyses of published single-cell datasets indeed showed significant enrichment of tumor-infiltrating T_FH_ cells following checkpoint blockade with anti-PD-1 agents, which suggested that T_FH_-like CD4^+^ T cells may also be important targets of anti-PD-1 therapies.

In summary, our study has revealed that T_FH_-like CD4^+^ T cells in the TME constitute a distinct functional subset that supports CD8^+^ CTL responses. Further studies will enable understanding of the mechanisms underlying generation and long-term maintenance of these cells and their functional significance in preventing relapse. Our findings suggest that eliciting a T_FH_ program in CD4^+^ T cells may be an important component of immunotherapies and vaccination approaches aimed to generate robust and durable CD8^+^ CTL and T_RM_ responses against neo-antigens or shared tumor antigens.

## MATERIALS AND METHODS

### Human subjects

Newly diagnosed, untreated patients with non-small cell lung cancer (**Supplementary Table 1**) referred to Southampton University Hospitals NHS Foundation Trust and Poole Hospital NHS Foundation trust, UK between 2014 and 2017 were prospectively recruited. Freshly resected tumor tissue and, where available, matched adjacent non-tumor tissue was obtained from patients with lung cancer following surgical resection.

### Mice

All mice were of C57BL6/J background and bred at LJI or purchased from the Jackson Laboratory. All mice inoculated with tumor were females and between 8-10 weeks of age at the beginning of experiments. Within each cage, age-matched mice were randomly allocated to control or experimental groups and the investigator was not blinded to the allocation during the experiment.

### Tumor model

Tumor cell lines tested negatively for mycoplasma infection and Plasmocin (InvivoGen) was used as a routine addition to culture media to prevent mycoplasma contamination. Mice were inoculated with 1.5x10^5^ B16F10-OVA cells subcutaneously into the right flank. For generation of T_FH_ cells for adoptive transfer to tumor-bearing mice, OT-II mice were injected intra-peritoneally with 100µg OVA (Invivogen) in alum (Invivogen) and spleens were harvested 1 week post-immunization for flow sorting OT-II CD4^+^ T_FH_ or T_EFF_ cells (**Supplementary Figure 3**). Identical numbers (6x10^5^ - 9x10^5^ cells) of T_FH_ or T_EFF_ cells were transferred adoptively into tumor-bearing mice by retro-orbital injection. For induction of T_FH_ cells in tumor-bearing mice, immunization was performed by footpad and tailbase injection of 10µg OVA in alum. Tumor size was monitored every other day, and tumor harvested at indicated time points for analysis of tumor-infiltrating lymphocytes. Tumor volume was calculated as ½ x D x d^2^, where D is the major axis and d is the minor axis, as described previously(69).

### Flow cytometry of fresh samples

Patient samples were processed as described previously (3). For sorting of fresh CD4^+^ TILs for transcriptomic analysis, cells were first incubated with FcR block (Miltenyi Biotec), then stained with a mixture of the following fluorescence-conjugated antibodies (BD Biosciences or BioLegend): anti-human CD45 (HI30), CD4 (RPA-T4), CD3 (SK7), CD8a (SK1), HLA-DR (L243), CD14 (MφP9), CD19 (HIB19) and CD20 (2H7) for 30 min at 4°C. Live/dead discrimination was performed by DAPI staining. CD4^+^ T cells were sorted into ice-cold TRIzol LS reagent (Ambion) using a BD FACSAria™ (BD Biosciences).

Murine samples –Tumor tissue was dispersed in 2ml of PBS, followed by incubation of samples in a shaker at 37°C for 15 min with DNase I (Sigma) and Liberase DL (Roche). The suspension was then diluted with MACS buffer and passed through a 70-µm cell strainer to generate a single cell suspension. Cells were first incubated with FcR block (clone 2.4G2, BD Biosciences), then with a mixture of antibodies for 30 min at 4°C. The following fluorescence-conjugated antibodies (BD Biosciences or BioLegend) were used in different panels for surface staining: anti-mouse Ctla-4 (UC10-4B9), Cd3 (145-2C11), Cd4 (RM4-5), Cd8 (53-6.7), Pd-1 (29F1.A12), Cd19 (6D5), Gitr (DTA-1), Cd45 (30-F11), Icos (C398.4A), Cxcr5 (L138D7), Tcrb, Samples were then sorted or fixed. Intracellular staining was performed using anti-mouse Ki67 (B56), FoxP3 (FJK-16s), Bcl6 (K112-91), GzmB (QA16A02) and FoxP3 transcription factor kit as per manufacturer’s protocol (eBioscience). Cell viability was determined using fixable viability dye (ThermoFisher). Samples were analyzed on a BD FACS Fortessa.

For sorting for adoptive transfer of OT-II CD4^+^ T_EFF_ cells or OT-II CD4^+^ T_FH_, splenocytes were first enriched for OT-II CD4^+^ T cells using EasySep™ mouse CD4^+^ T cell isolation kit (StemCell Technologies), incubated with FcR block and stained with the following fluorescence-conjugated antibodies: anti-mouse Cd45 (30-F11), Cd3 (145-2C11), Cd4 (RM4-5), Cd8 (53-6.7), Pd-1 (29F1.A12), Cd19 (6D5), Gitr (DTA-1), Icos (C398.4A), Cxcr5 (L138D7), Cd25 (PC61), Cd44 (IM7) and Cd62L (MEL-14). Cell viability was determined using fixable viability dye (ThermoFisher). Samples were sorted on a BD FACS Fusion (BD Biosciences).

### Flow cytometry of cryopreserved samples

For 10x single-cell transcriptomic analysis and phenotypic characterization, patient tumor and lung samples were first processed and cryopreserved in freezing media (50% complete RMPI (Fisherscientific), 40% human decomplemented AB serum, 10% DMSO (both Sigma). Cryopreserved samples were thawed, incubated with FcR block (Miltenyi Biotec), then stained with a combination of the following fluorescence-conjugated antibodies (BD Biosciences or BioLegend) for sorting: anti-human CD45 (HI30), CD3 (SK7), CD8A (SK1), CXCR5 (RF8B2), CD25 (MA251), CD127 (eBioRDR5), CD19/20 (HIB19/2H7), CD56 (HCD56) and CD4 (OKT4). Live/dead discrimination was performed using propidium iodide (PI). 1500 TILs from each of the three subsets, CD4^+^CXCR5^+^, CD4^+^CXCR5^−^CD25^−^ and CD4^+^CXCR5^−^CD25^+^CD127^−^ from tumor of each patient and 4500 CD4^+^ T cells (N-TILs) from adjacent uninvolved lung of each patient were sorted into 50% ice cold PBS, 50% FBS (Sigma) using a BD Aria-III (BD Biosciences).

For intracellular staining for the chemokine CXCL13 and granzyme B, TILs and N-TILs were incubated in RPMI 1640 medium (Life Technologies) containing brefeldin A (5ug/ul) for 3.5 hrs. TILs and N-TILs were stained using Zombie Aqua fixable viability kit (Biolegend), following which surface staining was performed with a mixture of fluorescence-conjugated antibodies (BD Biosciences or BioLegend): anti-human CD45 (HI30), CD3 (SK3), CD4 (RPA-T4), CD8 (SK1), CXCR5 (RF8B2), CD25 (2A3), CD127 (A019D5), PD-1 (EH12.2H7) for 30 min at 4°C. After fixation (BD Cytofix/Cytoperm) and permeabilization (BD Perm/Wash buffer), intracellular staining was performed with fluorescence-conjugated antibodies, anti-human CXCL13 (53610, R&D Systems) and GZMB (REA226, Miltenyl Biotec), for 30 min at 4°C. For intracellular staining for T regulatory cells, the following fluorescence-conjugated antibodies and buffers were used: anti-human Foxp3 (PCH101, ThermoFisher), CXCL13 (53610, ThermoFisher) and Foxp3 staining buffer kit (eBioscience). Samples were analyzed on a BD LSRII or ImageStreamX MkII imaging flow cytometer (Amnis, Seattle).

### ImageStream Analysis

Samples were processed as described above for flow cytometry. Images were acquired on a 2-camera ImageStreamX MkII imaging flow cytometer (Amnis, Seattle) at low speed with 40X objective and INSPIRE software version 200.1.620.0. The cytometer passed all ASSIST performance checks prior to image acquisition. BB515 (Ch02, 480-560 nm), PE (Ch03, 560-595 nm) PE-Dazzle594 (Ch04, 595-642 nm), PerCP-Cy5.5 (Ch05, 648-745 nm) and PE-Cy7 (Ch06, 745-780 nm) were excited at 488nm (40 mW). BV421 (Ch07, 435-505 nm) was excited at 405 nm (20 mW). APC (Ch11, 640-745 nm) and APC-Cy7 (Ch12, 745-780 nm) were excited at 642 nm (150 mW). The acquisition gate was set to include all single, in-focus, live, CD3^+^ events. Data was compensated and analyzed with IDEAS software version 6.2.64.0 using the default masks and feature set.

### Histology and immunohistochemistry

Deparaffinisation, rehydration, antigen retrieval and IHC staining was carried out using a Dako PT Link Autostainer. Antigen retrieval was performed using the EnVision FLEX Target Retrieval Solution, High pH (Agilent) for all antibodies. The primary antibodies used for IHC includes anti-CD103 (EPR4166(2); 1:500; Abcam), anti-CXCL13 (polyclonal; 1:100; ThermoFisher Scientific), anti-CD8 (C8/144B; pre-diluted; Agilent Dako), anti-CD4 (4B12; pre-diluted; Agilent Dako), anti-granzyme B (GrB-7; 1:50; Dako) and anti-PanCK (AE1/AE3; pre-diluted; Agilent Dako). Primary antibodies were detected using EnVision FLEX HRP (Agilent Dako) and either Rabbit or Mouse Link reagents (Agilent Dako) as appropriate. Chromogenic visualization was completed with either two washes for five minutes in DAB or one wash for thirty minutes in AEC and counterstained with hematoxylin. To analyze multiple markers on single sections, multiplexed IHC staining was performed as described previously(70). 4 micron tissue sections were stained with anti-PanCK antibody, visualized using DAB chromogenic substrate and scanned using a ZEISS Axio Scan.Z1 with a 20x air immersion objective. Each immune marker was then visualized using AEC chromogenic substrate and scanned. Between each staining iteration, antigen retrieval was performed along with removal of the labile AEC staining and denaturation of the preceding antibodies.

For each tissue section, regions within the tumor core (1 per section) or at the invasive margin (2 per section) were identified by a pathologist (GJT). These regions were exported as ome.tiff files and processed using Fiji image analysis software(71) as follows. The PanCK alone image was used as a reference for registering each iteration of staining, using the linear stack alignment with SIFT plugin. Color deconvolution for hematoxylin, DAB and AEC staining was performed using a customized vector matrix(72). 8bit deconvoluted images were then visually inspected to determine a pixel intensity threshold of positive staining for each marker and this value was subtracted from each image to remove non-specific staining. This color deconvolution approach resulted in DAB positive regions also being identified as AEC positive, therefore the PanCK alone image was used to generate a 0/255 pixel intensity binary “DAB mask”, which was then subtracted from each AEC image. Cell simulation and analysis was then performed using Tissue Studio image analysis software (Definiens). A machine learning classifier was trained to recognize epithelial and stromal regions using hematoxylin and PanCK staining. Cells were then identified by nucleus detection and cytoplasmic regions were simulated up to 5µm. CD4^+^CXCL13^+^ and CD8^+^CD103^+^ cells were then enumerated within the stromal regions of each image. This analysis was performed for 41 patients out of the total 45 patients in the cohort; due to insufficient sample, 4 patients were not analyzed.

### Bulk RNA sequencing

Total RNA was purified using a miRNAeasy micro kit (Qiagen, USA) and quantified as described previously (73) (on average, ∼8000 CD4^+^ T cells per sample were processed for RNA-seq analysis). Purified total RNA was amplified following the smart-seq2 protocol(73, 74). cDNA was purified using AMPure XP beads (1:1.1 ratio, Beckman Coulter). From this step, 1 ng of cDNA was used to prepare a standard Nextera XT sequencing library (Nextera XT DNA sample preparation kit and index kit, Illumina). Samples were sequenced using HiSeq2500 (Illumina) to obtain 50-bp single-end reads (**Supplementary Table 1**). Quality control steps were included to determine total RNA quality and quantity, optimal number of PCR pre-amplification cycles, and cDNA fragment size (73). Samples that failed quality control were eliminated from further downstream steps.

### 10x Single-cell RNA sequencing

Samples were processed using 10x v2 chemistry as per manufacturer’s recommendations; 11 and 12 cycles were used for cDNA amplification and library preparation respectively (57). Barcoded RNA was collected and processed following manufacturer recommendations, as described previously. Libraries were sequenced on a HiSeq2500 (Illumina) to obtain 100- and 32-bp paired-end reads using the following read length: read 1, 26 cycles; read 2, 98 cycles; and i7 index, 8 cycles (**Supplementary Table 1**).

### Bulk-RNA-seq analysis and iWGCNA

Bulk RNA-seq data were mapped against the hg19 reference using TopHat (75) (v1.4.1: --library-type fr-unstranded --no-coverage-search) and read counts were calculated using htseq-count -m union -s no -t exon -i gene_id (part of the HTSeq framework, version 0.7.1)) (76). Cutadapt (v1.3) was used to remove adapters.

WGCNA was completed using a R package WGCNA *(v1.61)* from the TPM data matrix generated from HTSeq-based read counts (28). Expressed genes with TPM >1 in at least 25% of the samples, were used in both CD4^+^ and CD8^+^ TIL data. In the integrated WGCNA approach, highly correlated genes from combined transcriptomes of patient-matched CD4^+^ TILS and CD8^+^ TILS were identified and summarized with a modular eigengene (ME) profile (28). Gene modules were generated using blockwiseModules function (parameters: checkMissingData = TRUE, power = 6, TOMType = “unsigned”, minModuleSize = 50, maxBlockSize = 25426, mergeCutHeight = 0.40) (**Supplementary Table 2**). Module 30, which represented the default ‘grey’ module generated by WGCNA for non-co-expressed genes, was excluded from further analysis. For each gene module, individual MEs were also calculated for CD4^+^ TIL-genes and CD8^+^ TIL-genes separately. As each module by definition is comprised of highly correlated genes, their combined expression may be usefully summarized by eigengene profiles, effectively the first principal component of a given module. A small number of eigengene profiles may therefore effectively ‘summarize’ the principle patterns within the cellular transcriptome with minimal loss of information. This dimensionality-reduction approach also facilitates correlation of ME with traits. Cell cycle signature was used to generate an eigengene vector from CD8^+^ TIL-genes, which was then used as a trait and correlated with MEs. Significance of correlation between this trait and MEs was assessed using linear regression with Bonferroni adjustment to correct for multiple testing.

To visualize co-expression network, we used the function *exportNetworkToCytoscape* at weighted = true, threshold = 0.05. A soft thresholding power was chosen based on the criterion of approximate scale-free topology. Networks were generated in Gephi (v0.92) (77, 78) using Fruchterman Reingold and Noverlap functions (**Supplementary Table 2**). The size and color were scaled according to the *Average Degree* as calculated in Gephi, while the edge width was scaled according to the WGCNA edge weight value.

The CD103 status of TILs was determined as previously described (3). GSEA determines whether an a priori defined ‘set’ of genes show statistically significant cumulative changes in gene expression between phenotypic subgroups using Kolmogorov-Smirnov statistic and was generated as previously described (3, 79). Genes used in the GSEA analysis are shown in **Supplementary Table 8**.

### Single-cell RNA-seq analysis

Raw 10x data was processed as previously described, merging multiple sequencing runs using *cellranger count* function in cell ranger, then merging multiple cell types with *cell ranger aggr (v2.0.2)*. The merged data was transferred to the R statistical environment for analysis using the package Seurat (v2.1) (57, 80). Only cells expressing more than 200 genes and genes expressed in at least 3 cells were included in the analysis. The data was then log-normalized and scaled per cell and variable genes were detected. Transcriptomic data from each cell was then further normalized by the number of UMI-detected and mitochondrial genes. A principal component analysis was then run on variable genes, and the first 6 principal components (PCs) were selected for TILs for further analyses based on the standard deviation of PCs, as determined by an elbow plot in Seurat. Cells were clustered using the *FindClusters* function from Seurat with default settings, resolution = 0.6. Clusters with less than 50 cells were excluded from analysis. Seurat software was used to identify cluster-specific differentially expressed gene sets (cutoff used is *q* < 0.05) (**Supplementary Table 3**).

Differential expression between two groups was determined by converting the data to CPM and analyzing group-specific differences using MAST (*q* < 0.05, v1.2.1) (57, 81, 82) (**Figure 2**; **Supplementary Table 4**). For differential expression between three groups (*CXCL13*-expressing single cells in CXCR5^+^ subset (*n* = 300), CXCR5^-^CD25^-^ subset (*n* = 319), CXCR5^-^CD25^+^CD127^-^ subset (*n* = 336)) (**Figure 3E**), pairwise comparisons were performed (**Supplementary Table 7**). A gene was considered significantly different (unique to a group), only if the gene was commonly positively enriched in every comparison for a singular group (57, 79). A gene was considered shared between two groups if the gene was commonly positively enriched in the two groups compared to the third group. For this analysis in **Figure 3E**, only genes that overlapped with those differentially expressed in *CXCL13*-expressing *versus CXCL13-*non-expressing cells (**Figure 2**; **Supplementary Table 4**) were used.

The mean CPM and percentage of cells expressing a transcript expressing cells was calculated with custom R scripts. Further visualizations of exported normalized data were generated using the Seurat package and custom R scripts. Average expression across a cell cluster was calculated using the *AverageExpression* function, and downsampling was achieved using the *SubsetData* function (both in Seurat).

The biological relevance of differentially expressed genes identified by MAST analysis was further investigated using the Ingenuity Pathways Analysis platform (**Supplementary Table 6**) as reported previously (3).

### Meta-analysis of published single-cell RNA-seq studies

From each of the 9 published single-cell RNA-seq datasets (6, 19–24, 26, 27), we extracted all the cells from the cluster(s) that were annotated as tumor-infiltrating CD4^+^ or CD8^+^ clusters. For each cell type (either CD4^+^ or CD8^+^), we then filtered out the cells with expression (>1 CPM for UMI data or >10 TPM for Smart-seq2 data) of the other cell type’s representative transcript (i.e., filter out cells from the CD4^+^ clusters with *CD8B* expression and from the CD8^+^ clusters with *CD4* expression) (**Supplementary Table 5**). We then integrated the remaining cells and their corresponding clusters from each study using the R package Seurat v3.0 for each cell type (83). For each dataset, cells that expressed less than 200 genes were considered outliers and discarded. FindIntegrationAnchors function was used to find correspondences across the different study datasets with default parameters (dimensionality = 1:30). IntegrateData function was used to generate a Seurat Object with an integrated and batch-corrected expression matrix. For each cell type, the 2000 most variable genes were used for clustering. We used the standard workflow from Seurat, scaling the integrated data, finding relevant components with PCA and visualizing the results with UMAP. The number of relevant components was determined from an elbow plot. UMAP dimensionality reduction and clustering were applied with the following parameters: 2000 genes, 15 principal components, resolution of 0.2, min.dis 0.05 and spread 2. *CXCL13*-expressing or *PDCD1*-expressing cells were identified based on criteria defined in **Supplementary Table 5**. Patient tumor samples with total CD4^+^ cells or total CD8^+^ cells <30 were excluded from further analysis to avoid sampling bias due to low cell numbers.

Cell cycle signature (**Figure 2G**) was derived by comparing the CD8^+^ cell cycle cluster with other CD8^+^ clusters. The CD8^+^ cells were then ranked based on expression of the cell cycle signature. In this rank order, the threshold for cell cycle signature-positive cells was marked at both the lower (5.3%) and upper (11.9%) bound of the percentage of cells that belong to the cell cycle cluster across different studies.

### Quantification and statistical analysis

Hypergeometric test using phyper function and p.adjust in R was used to calculate adjusted significance values for gene enrichment tests (**Figure 1E**). Comparison between two groups was assessed with Mann-Whitney test (**Figure 5A-C** and **Supplementary Figure 1D**) or two-tailed paired Student’s *t*-test (**Figure 2D**, 2H, 2J and **Supplementary Figure 2A**) using GraphPad Prism 7. Spearman correlation coefficient (*r* value) was calculated to assess significance of correlation between any two parameters of interest (**Figure 2G, 4C**).

### Contact for reagents and resource sharing

Please contact Dr. Vijayanand for reagents and resources generated in this study.

### Data availability

RNA sequencing data reported in this paper has been deposited in NCBI GEO (GSE118604).

### Study approval

The Southampton and South West Hampshire Research Ethics Board approved the study (Ref. 14/SC/0186), and written informed consent was obtained from all subjects (3). The Institutional Animal Care and Use Committees (IACUC) of the La Jolla institute for Immunology (LJI) approved all animal studies (Ref. AP00001126).

## AUTHOR CONTRIBUTIONS

A.-P.G., P.S.F., T.S.-E., C.H.O., F.A., and P.V. conceived the work, designed and analyzed experiments; D.S. and A.-P.G. performed micro-scaled RNA-seq experiments under the supervision of G.S. and P.V.; D.S., A.-P.G., B.P., A.M., C.R.S. performed data analysis under the supervision of F.A., C.H.O. and P.V.; A.-P.G., S.E. and A.W. performed experiments using mouse models; C.J.H. performed immunohistochemistry and data analysis under the supervision of G.J.T.; J.C., O.W. and E.M.G.-M. helped perform the cell isolations; S.J.C., A.A. and E.W. assisted in patient recruitment, obtaining consent and sample collection; S.C., S.B., A.-P.G. and P.V. designed experiments for T_FH_ induction in mice; A.-P.G., C.H.O., F.A. and P.V. wrote the manuscript. All authors read and approved the final text of the manuscript.

## ACKNOWLEDGMENTS

We thank M. Chamberlain, K. Amer, B. Johnson for assistance with recruitment of study subjects and processing of samples; J. Greenbaum at the LJI bioinformatics core for help with processing and analysis of sequencing data; Y. Altman at the Sanford Burnham Prebys Flow Cytometry Core (NCI grant P30 CA030199) and the James B. Pendleton Charitable Trust for support and access to Amnis Image Stream analysis.

Supported by the Wessex Clinical Research Network and the National Institute of Health Research, UK (sample collection), Cancer Research UK (digital pathology, accelerator award C11512/A20256 to C.H.O., G.J.T., P.V.), Faculty of Medicine of the University of Southampton (P.V., T.S.-E, C.H.O.), National Institutes of Health (K08 CA230164-01A1 to A.-P.G.) and the William K. Bowes Jr Foundation (P.V.).

