## Supplementary Figures for "CD4^+^ follicular helper-like T cells are key players in anti-tumor immunity"

The authors have declared that no conflict of interest exists. P.V receives research funding unrelated to this work from Pfizer.

Supplementary Figure 1

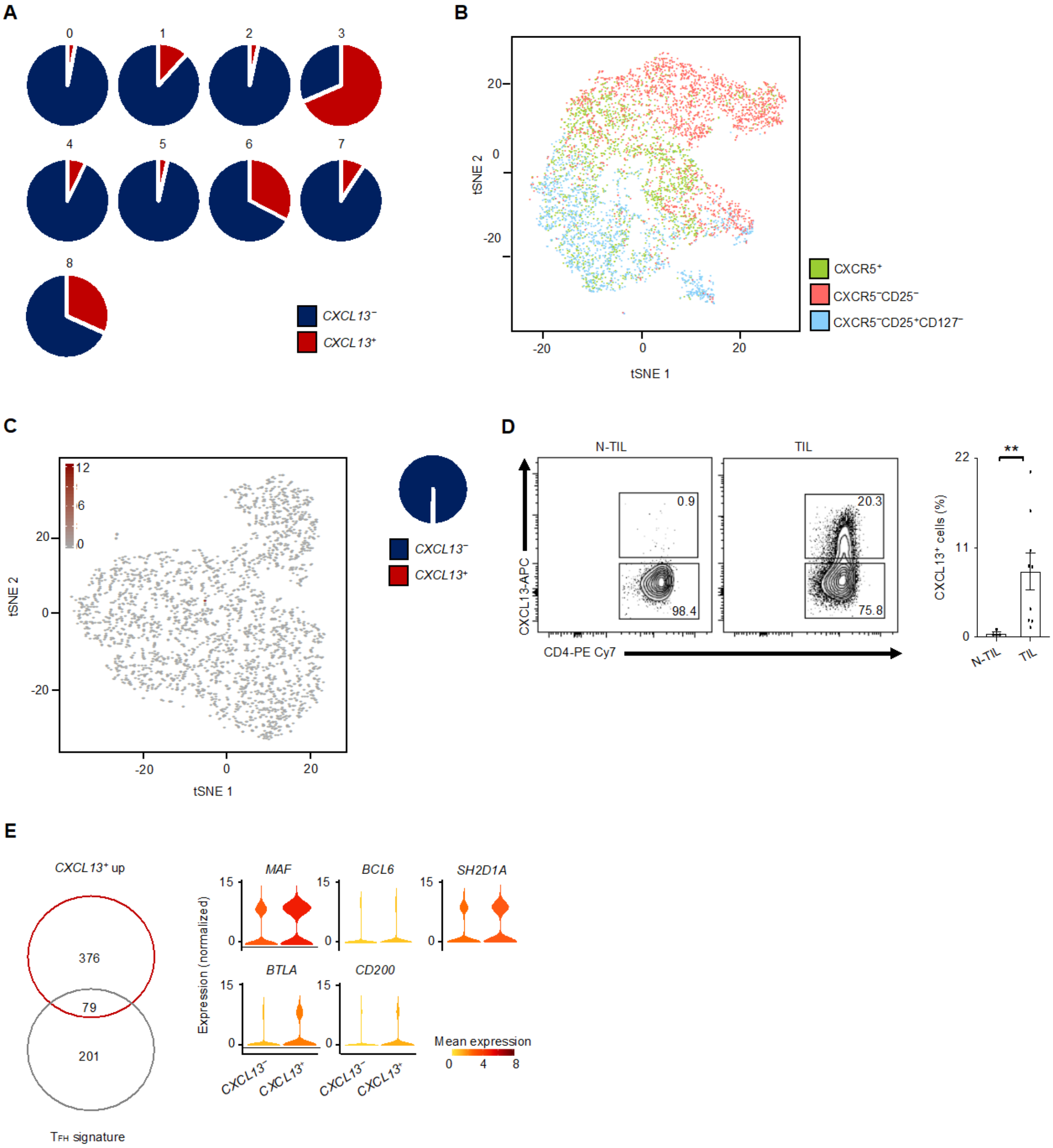


**Supplementary Figure 1. Single-cell transcriptomics reveal molecular profile of *CXCL13*-expressing tumor-infiltrating CD4⁺ T cells.** (**A**) Pie charts represent the proportion of *CXCL13*-expressing cells (key) in each cluster of CD4⁺ TILs. (**B**) tSNE visualization of cells in **Figure 2B**; each symbol represents a cell; color indicates the sorted subsets (key). (**C**) tSNE visualization (left) of 2783 single cell N-TIL transcriptomes; each symbol represents a cell; brown color indicates *CXCL13* expression (CPM). Pie chart (right) represents the proportion of *CXCL13*-expressing cells (key) among all N-TILs. (**D**) Flow-cytometric analysis shows the expression (left) and percentage (right) of CXCL13 in live, singlet-gated, $CD45^{+}CD3^{+}CD4^{+}$ T cells obtained from lung N-TILs (*n* = 4) or NSCLC TILs (*n* = 9) from patients with NSCLC. $^{**}P < 0.01$ (Mann-Whitney test). (**E**) Venn diagram (left) shows overlap of genes differentially expressed in *CXCL13*-non-expressing cells *versus* *CXCL13*-expressing cells and $T_{FH}$ signature genes. Violin plots (right) of expression of key $T_{FH}$ signature genes in *CXCL13*-expressing or *CXCL13*-non-expressing cells; shape represents the distribution of expression among cells and color represents expression ($\log_2$(CPM+1)).

**Supplementary Figure 2**

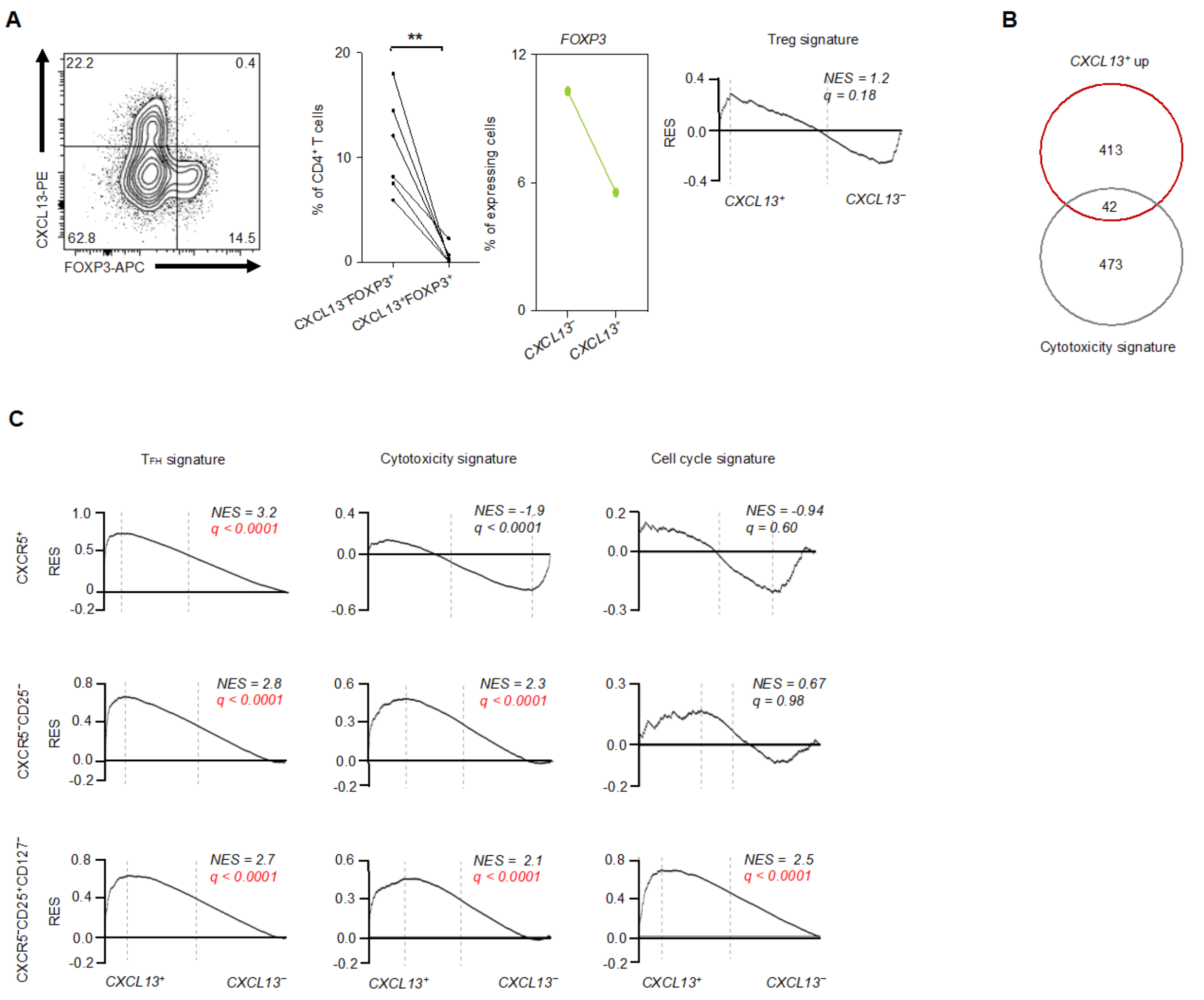


**Supplementary Figure 2**. **Highly functional $T_{FH}$-like CD4$^+$ T cells were CXCR5 negative.** (**A**) Flow-cytometric analysis shows the expression (left) and percentage (middle) of FOXP3 and CXCL13 in live, singlet-gated, CD45$^+$CD3$^+$CD4$^+$ TILs ($n$ = 6) from patients with NSCLC. **$P$ < 0.005 (two-tailed paired Student's $t$-test). Percentage (left margin) of *CXCL13*-non-expressing or *CXCL13*-expressing cells that express *FOXP3* (right). GSEA of Treg signature in the transcriptome of *CXCL13*-expressing *versus* CXCL13-non-expressing cells presented as in **Figure 1F** (far right). (**B**) Venn diagram shows overlap of genes differentially expressed in *CXCL13*-expressing cells *versus* non-expressing cells and cytotoxicity signature genes. (**C**) GSEA of various gene sets (above plots) in the transcriptome of *CXCL13*-expressing cells in the indicated subsets (left margin) *versus* all CXCL13-non-expressing cells from CD4$^+$ TILs presented as in **Figure 1F;** red font indicates significant enrichment in *CXCL13*-expressing cells.

**Supplementary Figure 3**

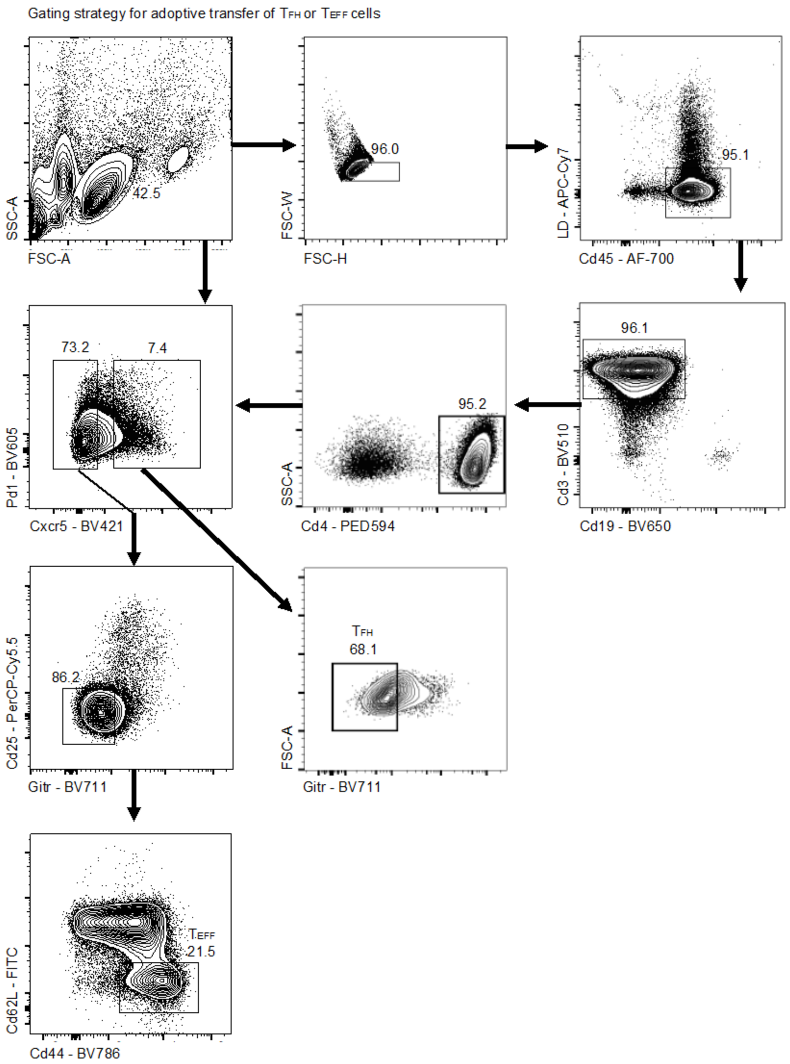


**Supplementary Figure 3. Adoptive transfer of OT-II CD4⁺ $T_{FH}$ cells or $T_{EFF}$ cells to tumor-bearing mice.** Gating strategy to sort OT-II CD4⁺ $T_{FH}$ (Cd19⁻Cd45⁺Cd3⁺Cd4⁺Cxcr5⁺Gitr⁻) or $T_{EFF}$ (Cd19⁻Cd45⁺Cd3⁺Cd4⁺Cxcr5⁻Gitr⁻Cd25⁻Cd62L⁻Cd44⁺) cells from spleens of OT-II mice following immunization with OVA in alum, shown are representative FACS plots.

**Supplementary Figure 4**

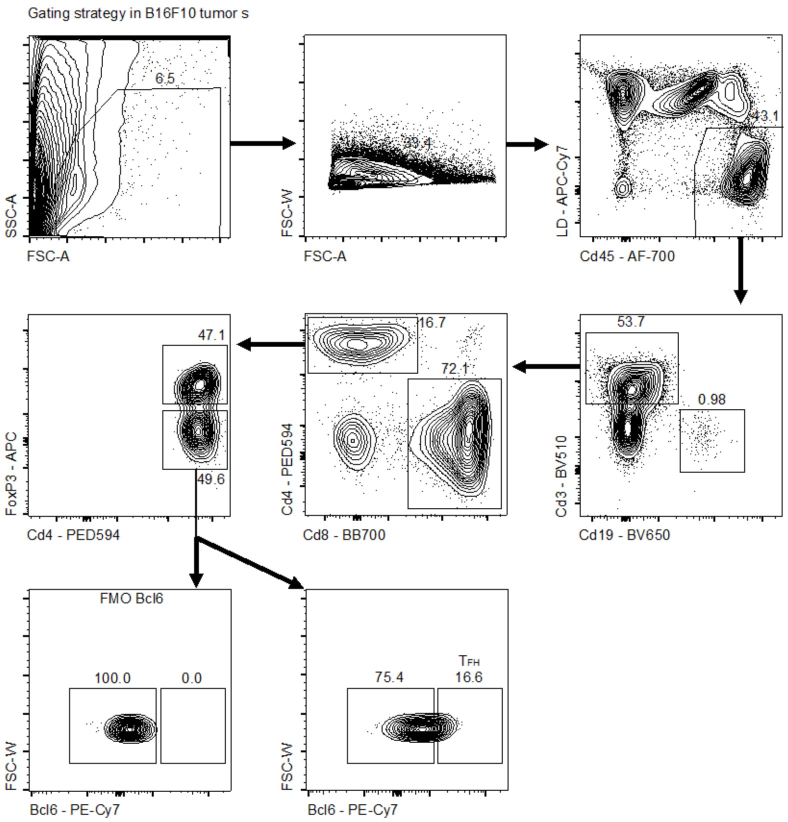


**Supplementary Figure 4. Characterization of CD4$^+$ T$_{FH}$ cells within murine tumors.** Gating strategy to identify tumor-infiltrating T$_{FH}$ (Cd19$^-$Cd45$^+$Cd3$^+$Cd4$^+$FoxP3$^-$Bcl6$^+$) cells in B16F10-OVA-inoculated mice at d21, shown are representative FACS plots.
